# Silicon Nitride Induces Osteoconduction Via Activated Mitochondrial Oxidative Phosphorylation and Neovascularization

**DOI:** 10.1101/2024.07.09.602787

**Authors:** Wayne Gonzales, Ronit Khade, Takeru Kondo, Satomi Arimoto, Taro Inagaki, Akishige Hokugo, Karolina Elżbieta Kaczor-Urbanowicz, Bryan McEntire, Ryan Bock, Takahiro Ogawa, Giuseppe Pezzotti, Ichiro Nishimura

## Abstract

Silicon nitride (Si_3_N_4_: SiN) is a thermodynamically stable ceramic material with excellent mechanical properties, and wear and corrosion resistance for industrial applications. SiN has been proposed for orthopedic and dental implant applications owing to its enhanced osteoconduction. However, the biological mechanisms underlying SiN-induced bone formation have not been fully elucidated. In this study, SiN significantly increased *in vitro* mineralization of human bone marrow mesenchymal stromal cells (BM-MSC) and *in vivo* peri-implant bone volume in mouse femurs compared to conventionally used titanium (Ti) implants. RNA sequencing of BM-MSC cultured on SiN disks revealed that functional gene clusters associated with mitochondrial oxidative phosphorylation were significantly elevated. SiN in an aqueous solution has been shown to release ammonium/ammonia, which may provide a source for glutamine-dependent energy production, and BM-MSC upregulated the expression of a key enzyme, glutamate-ammonia ligase under osteogenic conditions. Additionally, SiN increased the expression of functional gene clusters involved in vascular formation. The upregulation of *HIF1a in vitro* and increased VEGFR3-positive blanching vascular structures *in vivo* implied that SiN induced neovascularization. This study revealed an important mechanism through which SiN stimulated osteoconduction by unique glutamine-driven mitochondrial oxidative phosphorylation and established oxygen and nutrient supply by neovascularization, leading to stable osseointegration.

## 1. Introduction

Aging societies worldwide require the continuous productivity of the senior population, which requires the repair, regeneration, or replacement of diseased body parts. For example, the incidences of osteoarthritis, rheumatoid arthritis, and traumatic arthritis increase with age [1, 2], reflecting the recent increase in the utilization of prosthetic hip and knee surgeries among US Medicare beneficiaries [3]. While the use of dental implants has increased across different age groups, those aged 60 or older receive a larger number [4]. This treatment trend is consistent with increased tooth loss with advancing age. Approximately 17% of seniors 65 and older in the US have no remaining teeth (https://www.cdc.gov/oralhealth/publications/OHSR-2019-index.html).

Cementless prosthetic hip and knee implants and dental implants improve mechanical stability in the host bone bed through osseointegration [5, 6]. Osseointegration was initially thought to be induced on the surface of bioinert materials such as titanium (Ti) allowing bone wound healing at the surgical site without any harmful reactions or fibrosis. However, bone responses can be modulated on surface-modified Ti materials [7, 8] and zirconia (Zr) ceramic materials [9, 10], suggesting a novel opportunity to identify new biomaterials that can improve implant osseointegration for deteriorated host bone tissue of aging patients.

All load-bearing implant materials require mechanical stability. Silicon nitride (Si_3_N_4_; SiN) is an industrial ceramic material with outstanding thermodynamic stability used to manufacture highly durable components in the automotive, aviation, and aerospace industries. SiN exhibits mechanical strength, toughness, as well as wear and corrosion resistance, and is suitable for biomedical implants (**Table 1**) [11].

**Table 1.** Mechanical properties of silicon nitride (Si_3_N_4_: SiN), commercially pure titanium (Ti) and zirconia (Zr)

| Property | SiN |  | Ti |  | Zr |  | Units (S.I.) |
| --- | --- | --- | --- | --- | --- | --- | --- |
|  | Min | Max | Min | Max | Min | Max |  |
| Atomic Volume (average) | 0.0058 | 0.006 | 0.01 | 0.011 | 0.02 | 0.021 | m<sup>3</sup>/kmol |
| Density | 2.37 | 3.25 | 4.505 | 4.515 | 5 | 6.15 | Mg/m<sup>3</sup> |
| Energy Content | 150 | 200 | 750 | 1250 | 200 | 300 | MJ/kg |
| Bulk Modulus | 120 | 241 | 111 | 135 | 72.3 | 212 | GPa |
| Compressive Strength | 524 | 5500 | 130 | 170 | 1200 | 5200 | MPa |
| Ductility | 0.00031 | 0.00169 | 0.25 | 0.4 | 0.00066 | 0.0035 |  |
| Elastic Limit | 60 | 525 | 172 | 240 | 115 | 711 | MPa |
| Endurance Limit | 44 | 470 | 176 | 223 | 107 | 640 | MPa |
| Fracture Toughness | 1.8 | 6.5 | 55 | 60 | 1 | 8 | MPa.m<sup>1/2</sup> |
| Hardness | 8000 | 305000 | 1150 | 1250 | 5500 | 15750 | MPa |
| Loss Coefficient | 2.00E-05 | 5.00E-05 | 0.002 | 0.003 | 0.0005 | 0.001 |  |
| Modulus of Rupture | 181 | 1050 | 130 | 170 | 177 | 1000 | MPa |
| Poisson's Ratio | 0.23 | 0.28 | 0.35 | 0.37 | 0.22 | 0.32 |  |
| Shear Modulus | 65.3 | 127 | 36 | 39 | 53.4 | 86.4 | GPa |
| Tensile Strength | 60 | 525 | 240 | 360 | 115 | 711 | MPa |
| Young's Modulus | 166 | 297 | 100 | 105 | 100 | 250 | GPa |

SiNs exhibit biocompatible and osteoconductive properties. SiN disks [12], SiN-coated Ti [13], polyetherketoneketones [14], and polyetheretherketone [15, 16] increase *in vitro* bone formation of various osteogenic cells over conventional implant materials. These studies suggest that SiN enhances osteoconductivity; however, the underlying biological mechanisms are not fully understood.

Therefore, this study aimed to compare SiN with other commonly used implant materials using *in vitro* culture and *in vivo* mouse femur implant models. To elucidate the SiN-induced biological mechanisms, RNA sequencing was used to generate new hypotheses, which were validated in subsequent *in vitro* and *in vivo* experiments. The outcome of this study showed that the osteoconductive effect of SiN was manifested through novel mechanisms that activate glutamine-driven mitochondrial oxidative phosphorylation, which provided the energy necessary for new bone formation and induced neovascularization, ensuring oxygen and nutrient supply for further bone maturation.

## 2. Results

### 2.1. In vitro evaluation of SiN, Machined Ti (MTi), Acid-etched Ti (ATi), and Zirconia (Zr) disks (Figure 1A)

This study first compared SiN with currently used dental implant materials, such as Ti with machined (MTi) and acid-etched (ATi) surfaces, as well as ceramic-based zirconia (Zr). The surfaces of the ceramic SiN and Zr disks exhibited characteristic crystalline nanostructures, whereas the metallic MTi and ATi disks had anisotropic and isotropic surfaces, respectively (**Figure 1B**). Surface roughness measurements indicated that the SiN and Zr disks presented similar Sa, Sz, and Sdr values, whereas ATi and MTi disks showed the highest and lowest values, respectively (**Figure 1C**). The contact angle measurements were similar for the hydrophilic SiN, MTi, and Zr disk samples. The contact angle of ATi was found to be smaller than that of the other disk samples (**Figure 1D**) and this observation was consistent with published data that the increased surface roughness of Ti was generally associated with decreased contact angle measurements [17].

**Figure 1.**
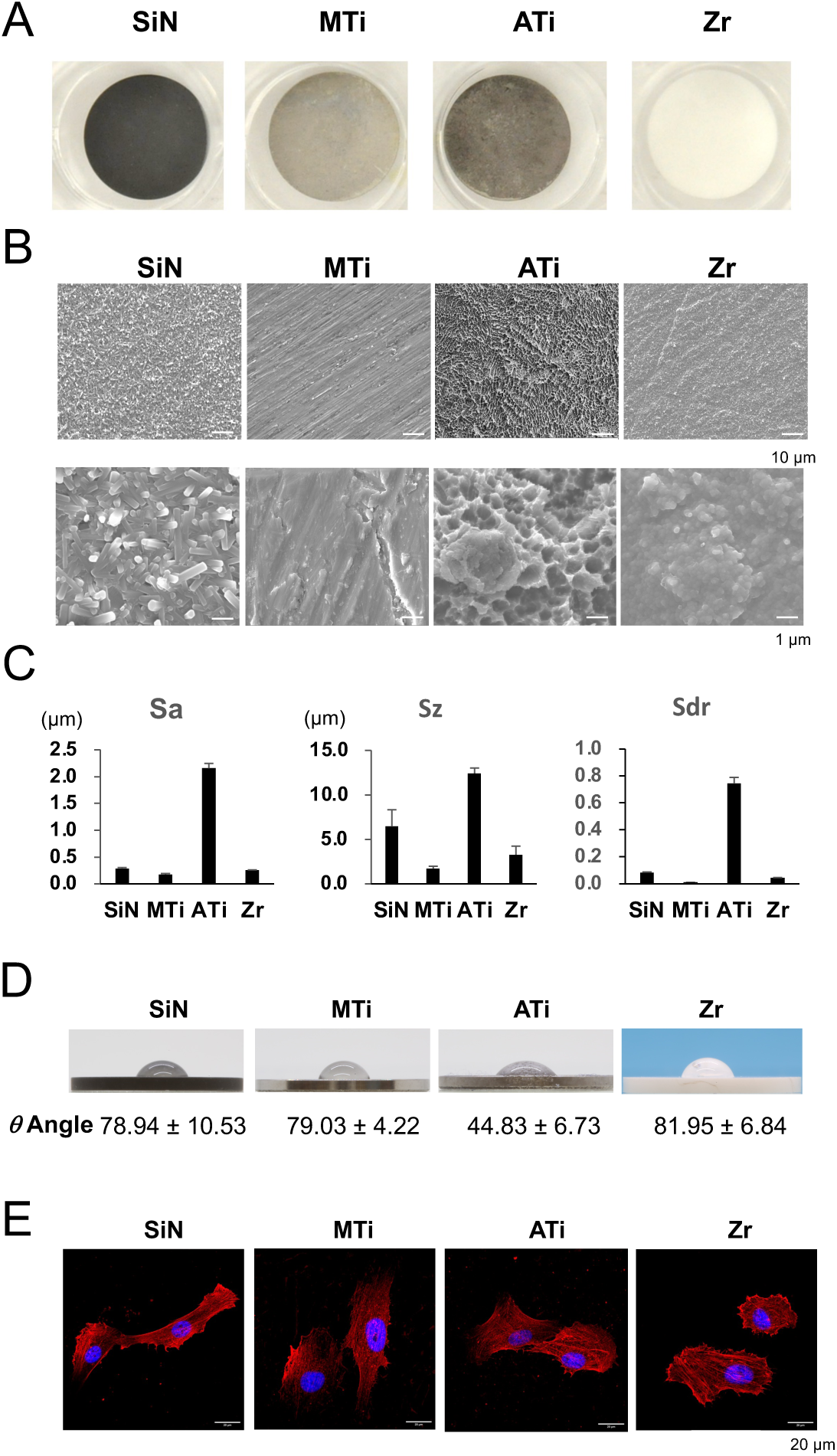
Characterization of Silicon Nitride (SiN), Machined and Acid-etched Titanium (MTi, ATi), and Zirconia (Zr) materials. **A.** Disks (12.7 mm diameter, 1 mm thick) were generated for *in vitro* study using SiN (Si_3_N_4_), MTi, ATi, and Zr. **B.** SEM surface characterization revealed the nanocrystalline structure of SiN and Zr ceramic materials. MTi and ATi materials showed anisotropic and isotropic surface structures, respectively. **C**. Surface topography characterization demonstrated micro-rough surface of ATi material (n=6 per group). **D**. Wettability test revealed that all materials were in the hydrophilic range. (n=6 per group). **E**. Human BM-MSC cultured on the materials were spread with developed cytoskeleton. No differences were observed.

Human bone marrow mesenchymal stromal cells (BM-MSC) cultured on SiN, MTi, ATi, and Zr disks attached to the material surface and developed stress fibers (**Figure 1E**). The cell growth measured by the Wst-1 assay suggested that the SiN disk appeared to slow down the cell proliferation rate, although statistical analysis did not show any major differences among the disk groups (**Figure 2A**). An inverse relationship between cell proliferation and differentiation has been previously demonstrated [18]. A striking observation in the present study was the robust increase in alizarin red S staining of BM-MSC cultured on SiN disks for 21 days, which was significantly higher than that in the other groups (**Figure 2B**).

**Figure 2.**
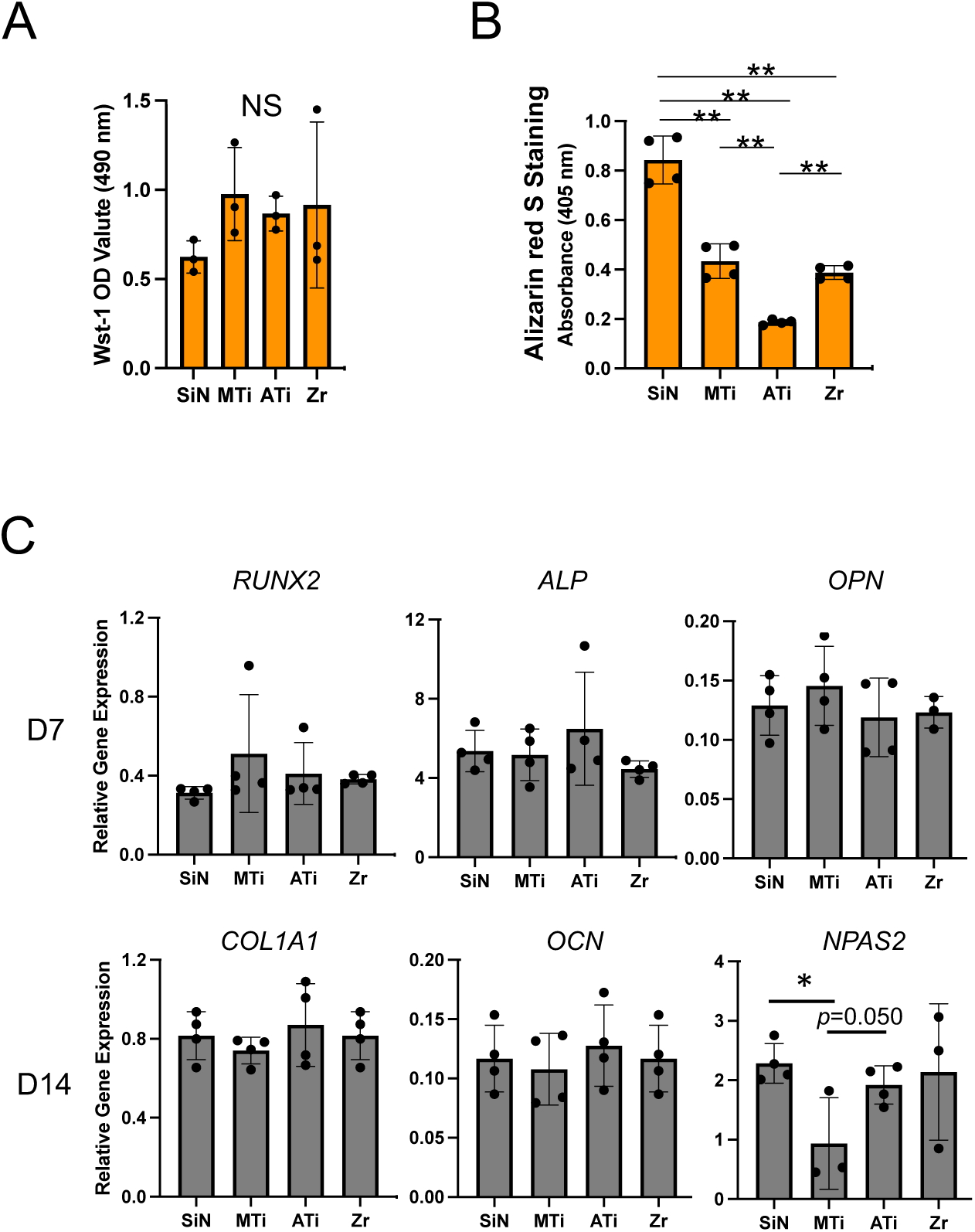
Behaviors of BM-MSC exposed to SiN, MTi, ATi and Zr disks. **A**. Wst-1 assay demonstrated that these materials did not affect the cell viability and proliferation of BM-MSC (n=3 per group). **B**. *In vitro* mineralization measured by alizarin red S staining increased in the SiN group (n=4 per group). **C**. RT-PCR analysis (n = 3∼4 per group) did not identify the effect of materials on the expression of osteogenic differentiation-related genes. However, a core circadian rhythm gene, *Npas2*, was upregulated by SiN and ATi as compared to MTi, albeit the statistical significance was reached only between SiN and MTi.

### 2.2. Gene expression analysis

Next, we investigated the effects of SiNs on osteogenic differentiation. Total RNA samples from BM-MSC cultured on SiN, MTi, ATi, and Zr disks for 7 and 14 days were isolated for RT-PCR. The steady-state expression of osteogenic differentiation-related genes such as Runx family transcription factor 2 (*RUNX2*), alkaline phosphate (*ALP*), osteopontin (*OPN*), alpha 1 type I collagen (*COL1A1*), and osteocalcin (*OCN*) did not show significant differences among the SiN, MTi, ATi, and Zr groups (**Figure 2C**), suggesting that the BM-MSC used in this study exhibited fate-specific differentiation into osteoblasts. However, these materials did not affect this process. A circadian rhythm gene, neuronal PAS domain 2 (*Npas2*), was previously reported in the peri-implant bone associated with Ti materials with micro-rough surfaces [19, 20], and was determined to be indispensable for achieving increased osseointegration of the surface- roughened Ti implant [21]. As expected, BM-MSC on the ATi disk showed an increase of the *Npas2* expression over the MTi disk (p=0.05 in the Turkey test and p<0.05 between ATi and MTi group by Student’s t test). In this study, BM-MSC cultured on SiN disks also demonstrated increased *Npas2* expression compared to MTi disks (p<0.05, Tukey’s test) (**Figure 2C**).

### 2.3. RNA sequencing and validations

To further investigate the molecular and cellular mechanisms underlying the increased *in vitro* mineralization induced by SiN disks, total RNA samples from the SiN, MTi, and ATi groups were subjected to RNA sequencing. Transcriptome data were analyzed for differentially expressed and coregulated genes. Gene ontology (Molecular Function and Cellular Component) as well as KEGG (Kyoto Encyclopedia of Genes and Genomes) pathways indicated that SiN disproportionately enriched the gene clusters involved in mitochondrial oxidative stress and oxidative phosphorylation (**Figure 3A**).

**Figure 3.**
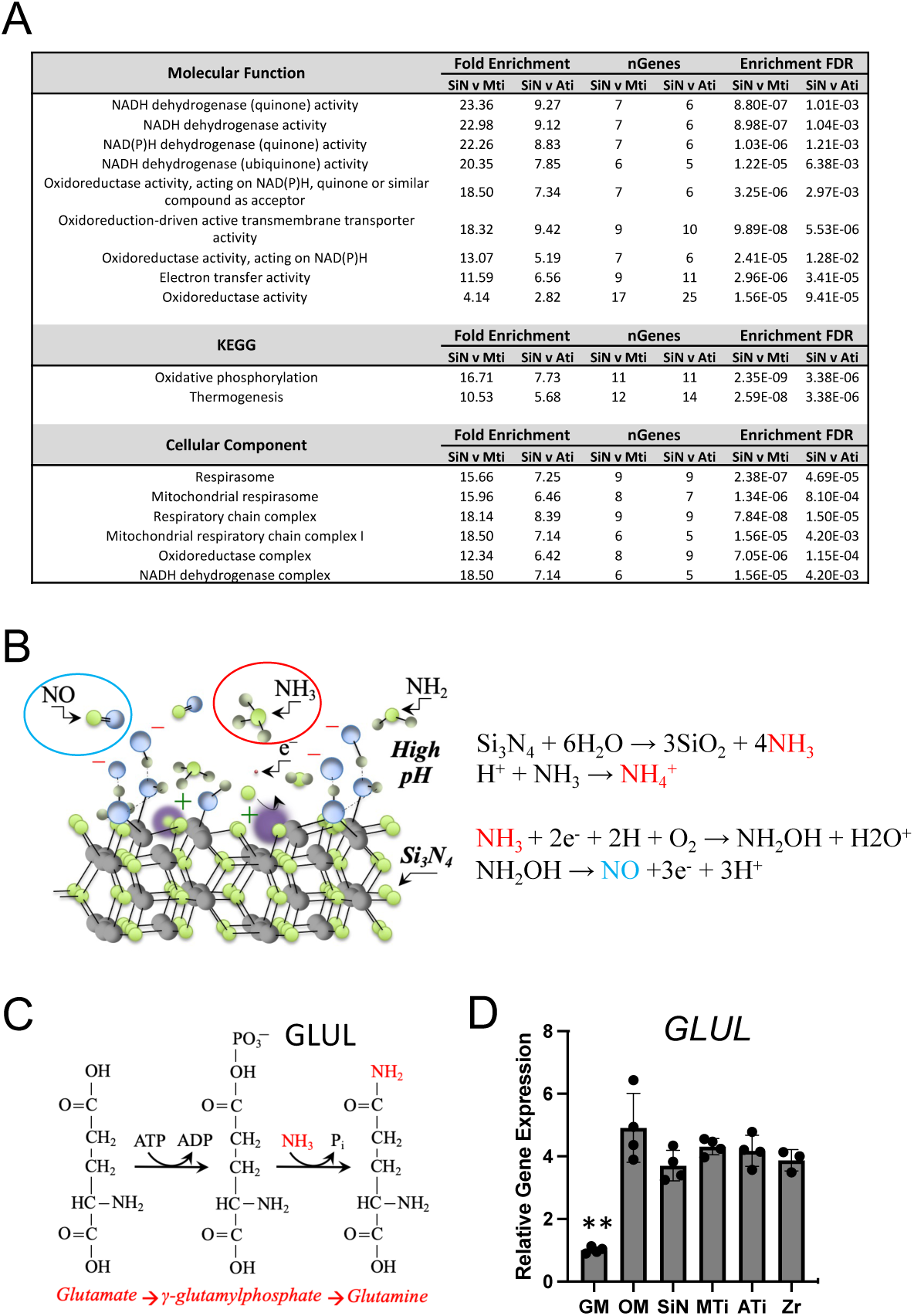
RNA sequencing data analysis revealed enrichment of mitochondria oxidative phosphorylation by SiN. **A**. Molecular Function and Cellular Component gene ontogeny as well as KEGG pathway analyses depicted that the functional enrichment of mitochondrial oxidative phosphorylation pathways in BM-MSC exposed to SiN. **B**. Hypothetical diagram of ammonium/ammonia released from SiN material, which may be integrated in the glutamine-driven mitochondrial oxidative phosphorylation. In addition, SiN-derived ammonia is thought to processed to generate nitric oxide (NO). **C**. Chemical diagram processing glutamate to glutamine by glutamine-ammonia ligase (GLUL). **D**. RT-PCR revealed the upregulation of *GLUL* by osteogenic media but it was not influenced by biomaterials.

It has been reported that SiN in aqueous solution releases nitrogen ions that interact with water molecules to form ammonium/ammonia [22, 23]. Furthermore, Raman spectroscopy demonstrated that the SiN-derived ammonia formed nitric oxide (NO) owing to the presence of free electrons formed from the splitting of water molecules [22, 24] (**Figure 3B**). The formation of NO has been experimentally proven using *in situ* fluorescence microscopy [20].

Thus, we hypothesized that SiN-derived ammonium/ammonia is a source of glutamine-driven mitochondrial oxidative phosphorylation [25] (**Figure 3C**). Among nutrients such as glucose and fatty acid [26], glutamine provides an efficient substrate for mitochondrial oxidative phosphorylation to generate energy in the form of adenosine triphosphate (ATP) [25]. Pathological depletion of glutamine from the bone marrow impairs the osteogenic differentiation of BM-MSC [27]. Glutamine is synthesized intracellularly by glutamine synthetase (glutamate-ammonia ligase, GLUL) (**Figure 3C**) [28]. RT-PCR demonstrated that steady-state *GLUL* expression was significantly increased in BM-MSC cultured in osteogenic medium for 21 days; however, it was not modulated by the biomaterials (**Figure 3D**).

Next, the RNA sequence data of BM-MSC cultured on SiN, MTi, and ATi disks were analyzed using Biological Pathway gene ontogeny, which revealed that SiN stimulated “extracellular matrix organization” and “extracellular structure organization” genes (**Figure 4A**) and that type VI collagen genes were upregulated. Furthermore, a large proportion of the top 20 biological pathways was related to angiogenesis, including blood vessel morphogenesis and vascular development (**Figure 4A**). The signature gene in the angiogenic biological pathway cluster was hypoxia-inducible factor 1a (*HIF1a*). The significantly increased steady-state expression of *HIF1a* was determined via RT-PCR in BM-MSC cultured on SiN disks compared to those cultured on MTi and ATi disks (**Figure 4B**).

**Figure 4.**
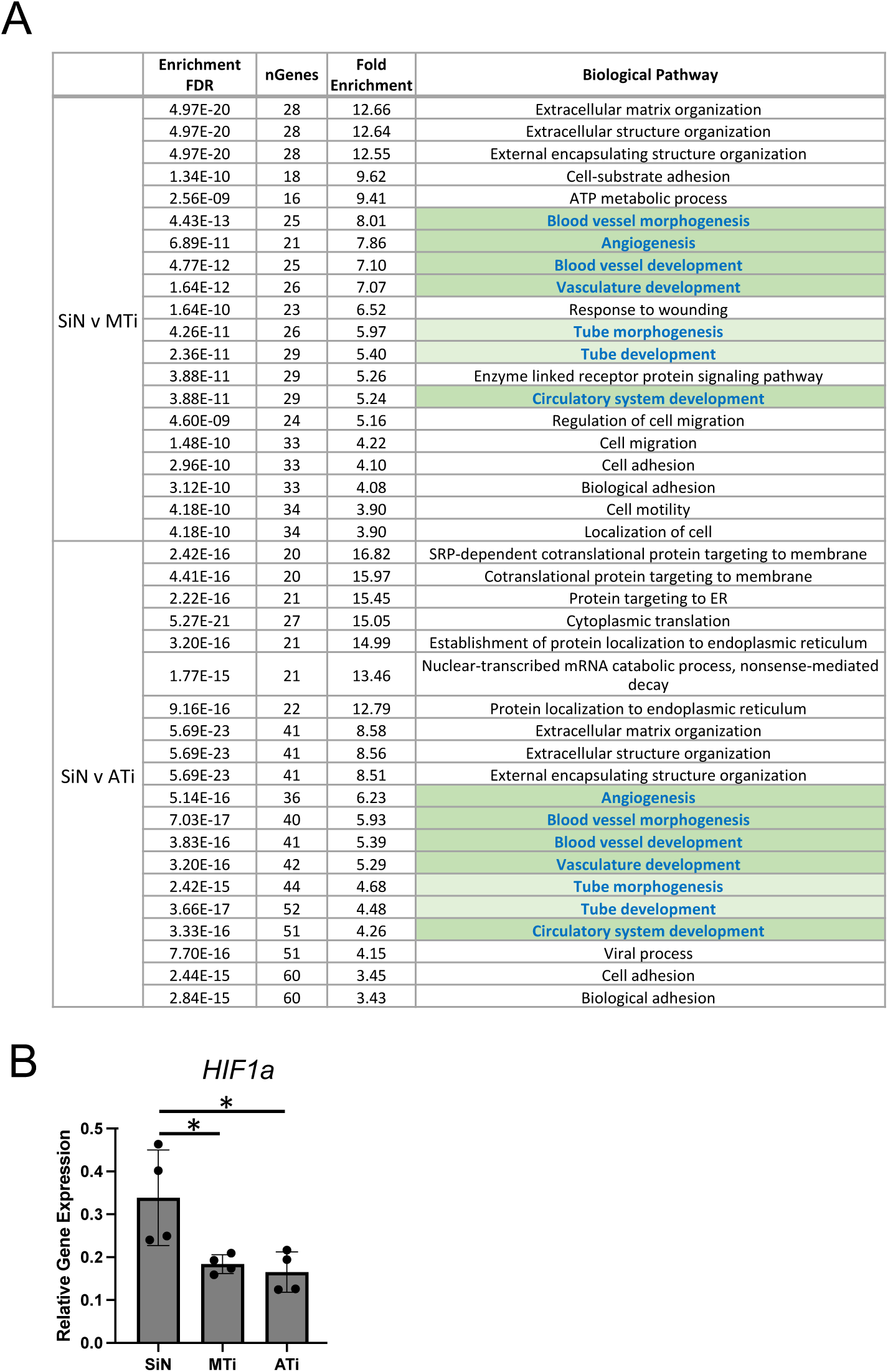
RNA sequencing data analysis by Biological Pathway. **A**. Top 20 Biological Pathway gene ontogeny included the increased gene clusters related to vascular formation (green shade) in the SiN group. **B**. Validation RT-PCR demonstrated the increased steady-state expression of the key gene, *HIF1a*, in the SiN group.

### 2.4. SiN, MTi, and ATi implants in the mouse femur model

Miniature implants (0.8 mm D and 6 mm L) were fabricated from SiN, MTi, and ATi (**Figure 5A**) and surgically placed in mouse femurs (**Figure 5B**), following previously reported protocols [29]. After 1 and 3 weeks of healing, implant osseointegration was measured using the implant push-in test after one and three weeks of healing (**Figure 5C**). The SiN and ATi implants resisted mechanical loading during the 1-week healing period, generating a push-in value, that is, the force required to dislodge the implant. The push-in value in the 3-week period increased and the ATi implant exhibited the largest push-in value, followed by SiN and MTi implants (**Figure 5D**).

**Figure 5.**
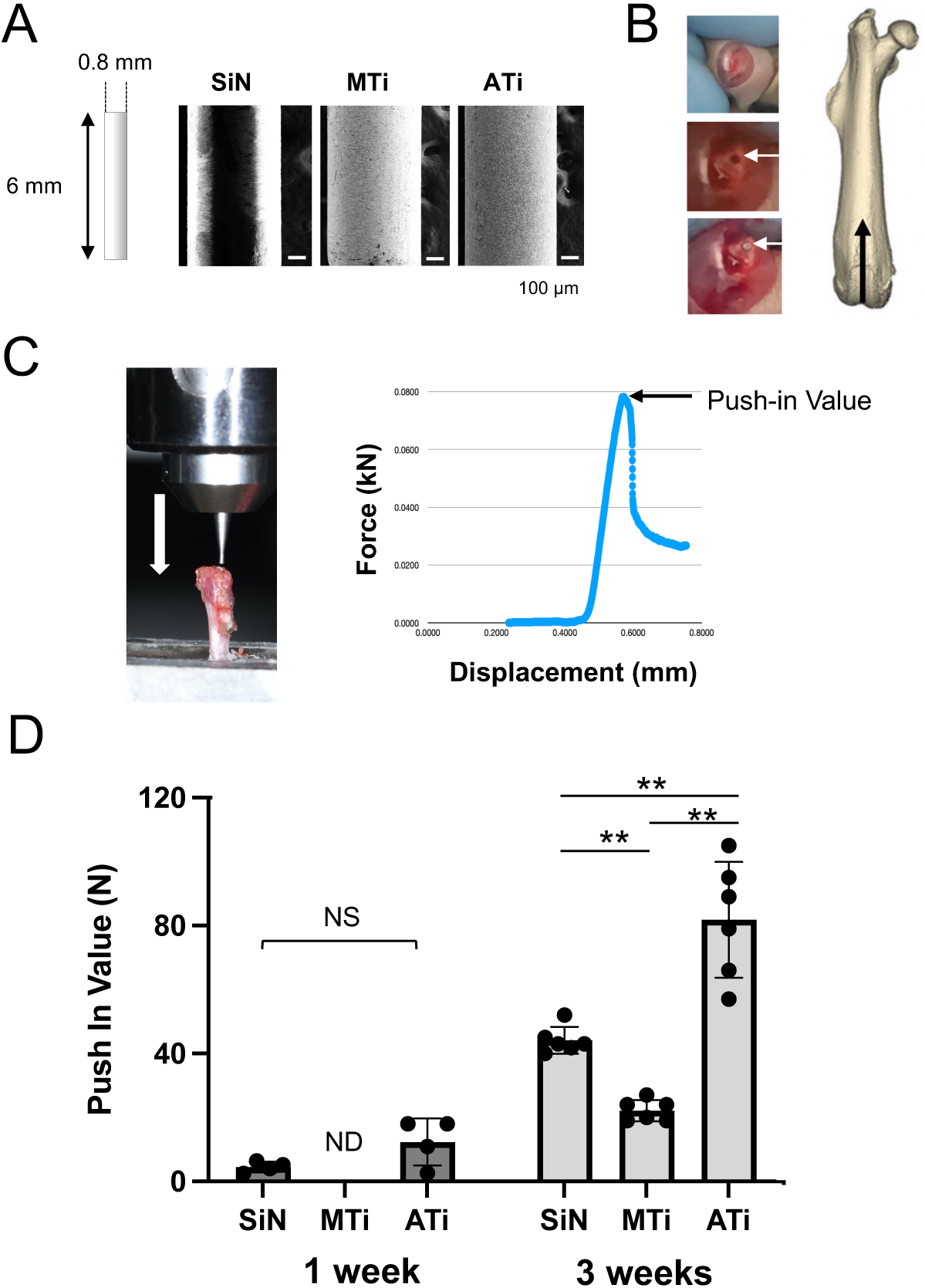
Osseointegration analysis of SiN, MTi, and ATi miniature implants in mouse femur. **A**. Miniature implants were fabricated for mouse femur. **B**. Surgical placement of miniature implant to mouse femur. **C**. Implant push-in test. The push-in value was determined at the break point of bone and implant bonding. **D**. Implant push-in value at 1 week (n=4 per group) and 3 weeks (n=6 per group) of healing time.

Mouse femur specimens containing miniature implants were fixed in 10% buffered formalin and subjected to micro-CT imaging. The 100 µm periphery from the surface of the implant embedded in the bone marrow was selected as the region of interest (ROI) (**Figure 6A**) and bone morphometry was measured. The peri-implant bone associated with the SiN implant exhibited a larger bone volume (BV/TV) and bone surface (BS/TV), smaller trabecular space (Tb.Sp), and more trabecular number (Tb.N) than those associated with the MTi and ATi implants (**Figure 6B**). Furthermore, the smallest structural model index (SIM) was observed in the SiN group, suggesting a more laminar peri-implant bone structure (**Figure 6B**).

**Figure 6.**
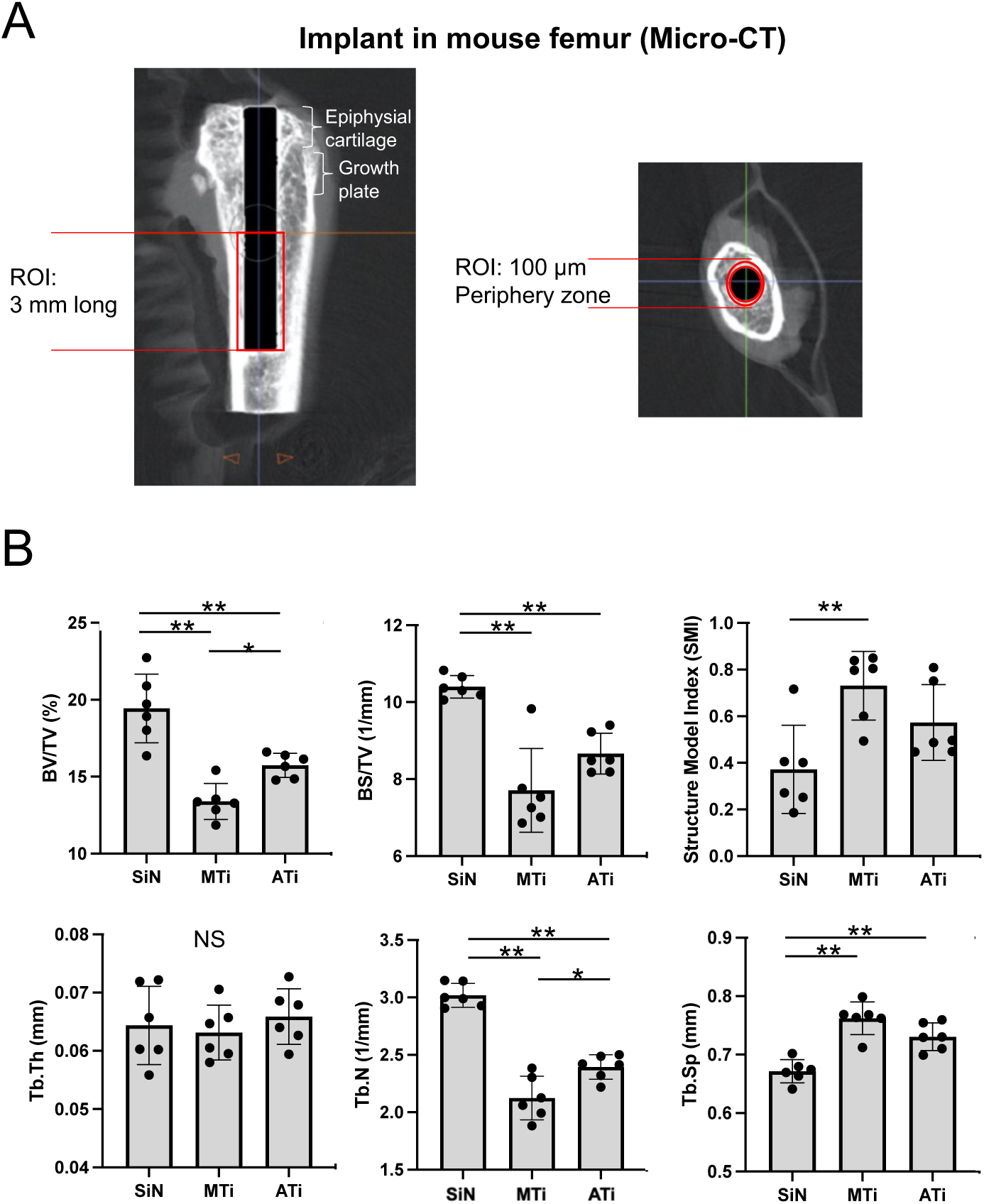
Micro-CT bone morphometry of peri-implant bone. **A**. A longitudinal section and a cross section micro-CT images depicting the peri-implant bone region of interest (ROI). SiN implant was radiolucent. **B**. Peri-implant bone morphometric measurement (n = 6 per group). Significant increase in the SiN peri-implant bone was suggested.

After micro-CT imaging, the mouse femur samples were decalcified and longitudinal histological sections were prepared after implant removal. Long and continuous peri-implant bones were observed in the SiN group, whereas peri-implant bones in the MTi and ATi groups were less frequent or fragmented (**Figure 7A**). Bone marrow around the MTi implant was associated with increased adipose tissue. The peri-implant bone surface forming the bone-to-implant contact (BIC) showed periodically spaced connective tissue structures in the SiN group (**Figure 7B**). In contrast, the BIC surfaces of the MTi and ATi implants were relatively smooth. A thin layer of nuclei was found to cover the peri-implant bone surface toward the MTi implant (**Figure 7B**), which was not included in the BIC ratio measurement. The BIC ratio was the largest in the SiN group (**Figure 7C**). The thickness of the peri-implant bone varied significantly; however, the mean thickness was widest in the SiN group (**Figure 7D**), supporting the micro-CT outcome.

**Figure 7.**
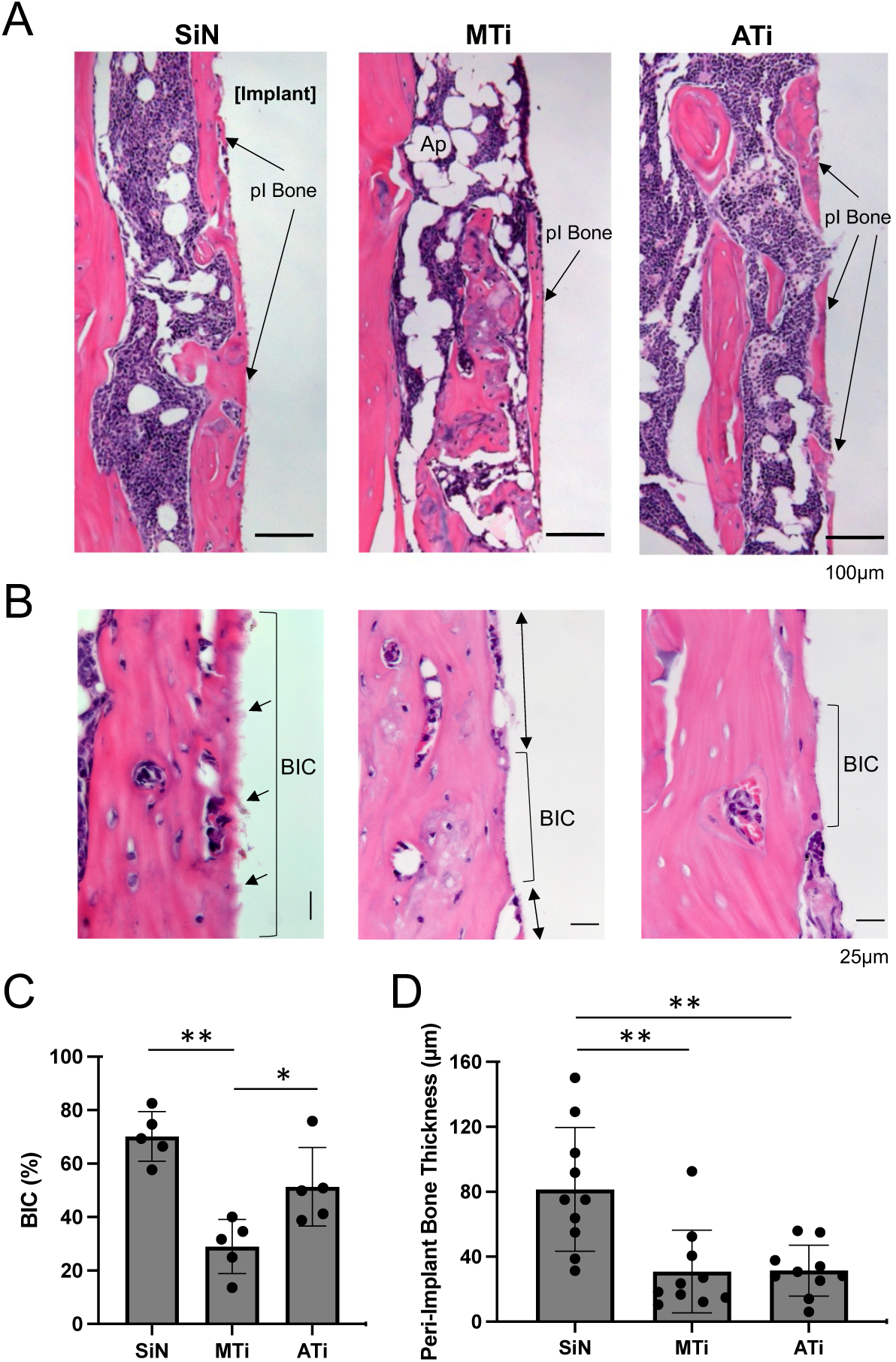
Histological evaluation of peri-implant bone. **A**. Peri-implant bone (pI Bone) associated with SiN implant was thick and well adopted. By contrast, peri-implant bone of MTi was thin and small and of ATi was fragmented. The bone marrow around MTi implant showed increased adipose tissue (Ap). **B**. High magnification view showed periodic connective tissue fibers (arrows) extended in the bone-to-implant contact (BIC) of SiN implant. There was a layer of cells between the peri-implant bone and MTi implant. **C**. BIC rate of SiN Implant and ATi implant was larger than MTi implant (n=5 per group). **D**. SiN associated peri-implant bone was thicker (wider) than those of MTi and ATi implants (10 measurement sites per group).

### 2.5. SiN induced angiogenesis

Immunohistochemistry was performed to validate the RNA sequencing data. The vascular endothelial cell marker CD31 was only weakly positive in the bone marrow tissue around the SiN implant and was not clearly observed in the MTi and ATi groups (**Figure 8A**). In contrast, microvasculature-like structures positive for vascular endothelial growth factor (VEGF) receptor 3 (VEGFR3) were clearly observed in the SiN group, and to some extent, in the ATi group (**Figure 8B**). The microvascular structures of the MTi group were negative for VEGFR3 expression. A greater number of VEGFR3-positive microvasculature structures were confirmed in the SiN group than in the MTi and ATi groups (**Figure 8C**).

**Figure 8.**
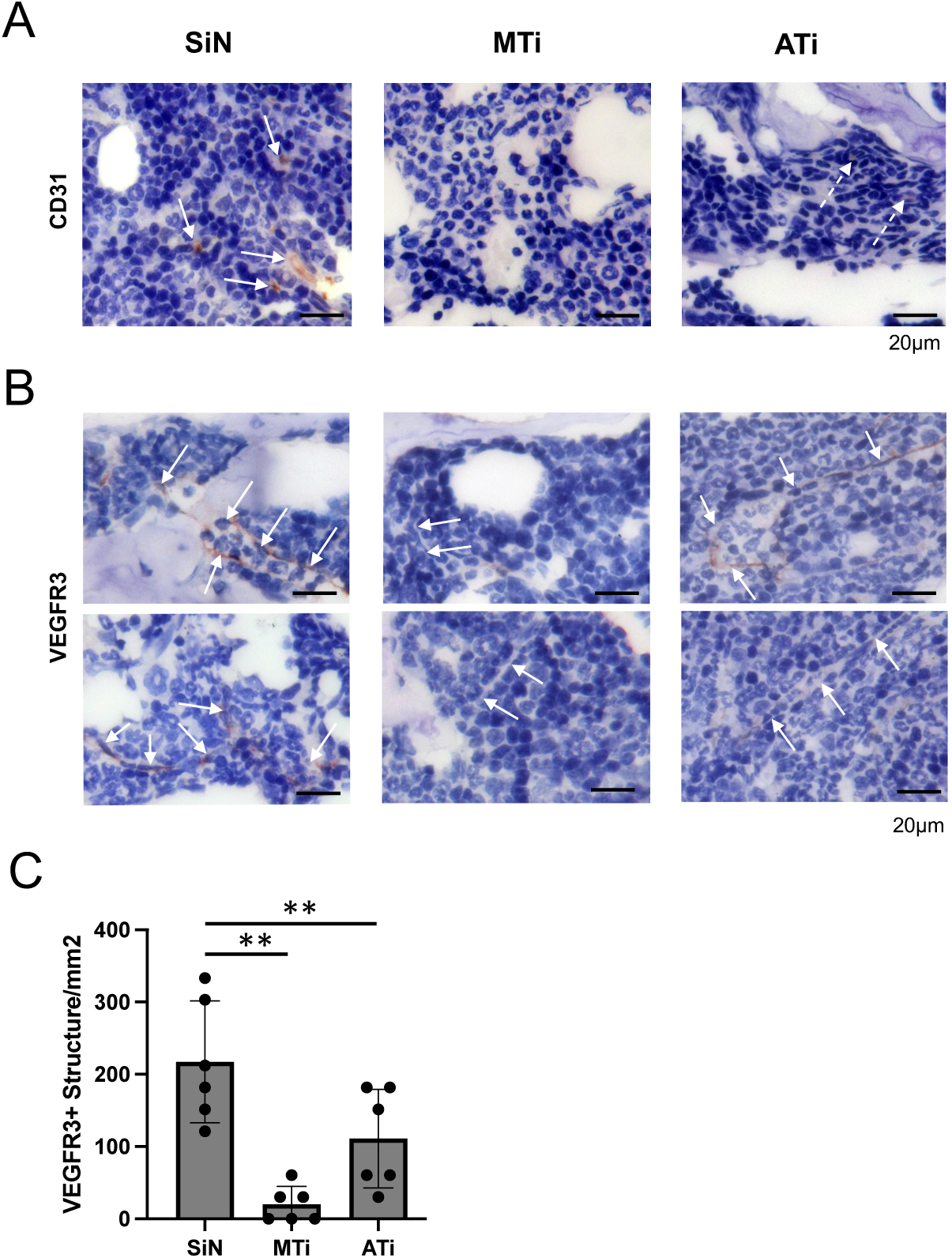
Increased CD31^dim^VEGFR3^+^ vascular structures in bone marrow around SiN implant. **A**. Immunohistochemistry with anti-CD31 antibody depicted faint staining (while arrows). **B**. Immunohistochemistry with anti-VEGFR3 showed strong staining of vascular-like structures in peri-implant bone marrow of SiN implant (white arrows). There were vascular-like structures in peri-implant bone marrow of MTi implant, which were negative with VEGFR3 staining (white arrows). In peri-implant bone marrow of ATi implant, there were VEGFR3 positive vascular-like structures (white arrows). **C**. VEGFR3 positive vascular-like structure was found more densely in peri-implant bone marrow of SiN implant.

## 3. Discussion

The present study revealed unique osteoconductive mechanisms that contributed to the osseointegration of SiN implants. Unlike osteoinduction, which involves the recruitment and differentiation of immature cells to pre-osteoblasts, osteoconduction is defined as the stimulation of fate-committed pre-osteoblasts toward bone formation within the bone microenvironment [30]. The present study demonstrated a robust increase in *in vitro* mineralization of human BM-MSC cultured on SiN disks (**Figure 2**) and *in vivo* bone generation around SiN implants in the mouse femur bone marrow (**Figures 5-7**).

Biomaterial-induced bone formation was characterized by the upregulation of selected osteogenic differentiation genes by RT-PCR. However, DNA microarray and more recent RNA sequencing data have not associated the expression of these genes with osteoconduction [31, 32]. Bone formation is an energy-consuming biological process and osteoblasts are metabolically active requiring high levels of energy from ATP [33, 34]. Mitochondria are cytosolic organelles that take up glucose-derived molecules, fatty acids, amino acids, and other fuels for oxidative phosphorylation to generate ATP. Targeted deletion of the core subunit of mitochondrial complex I in mouse skeletal progenitor cells results in deficient bone embryogenesis and spontaneous bone fractures [35]. Cyclophilin D-knockout mice, a genetic model of high oxidative metabolism, show greater osseointegration at spinal fusion sites [36]. The role of implant materials in regulating mitochondrial activity in the peri-implant microenvironment has recently gained attention [37]. In the present study, functional enrichment analysis of the RNA sequencing data revealed that SiN stimulated the upregulation of mitochondrial oxidative phosphorylation-related gene clusters compared to MTi and ATi (**Figure 3**), whereas osteogenic differentiation-related genes were not modulated by these materials (**Figure 2**).

When SiN (Si_3_N_4_) is exposed to aqueous solutions, the Si-N covalent bond is cleaved and nitrogen is leached; it reacts with water molecules to form ammonium (NH_4_^-^) and ammonia (NH_3_). Pezzotti (2019) reported that the ratio between ammonium and ammonia released from SiN was sensitive to fluid pH and predominantly ammonium was released under physiological conditions (pH7.4) at the concentration range of 0.01 mMol/dm^3^ [22]. While the SiN-released ammonia content is too low to be toxic, ammonium may positively affect the bone marrow cells surrounding the SiN implant. Liu et al. (2022) reported that BM-MSC, which highly express GLUL in the bone marrow, can convert ammonium and glutamate to glutamine and increase proliferation and osteogenic differentiation rates [38]. Glutamine is a major fuel of nitrogen and carbon for mitochondria oxidative metabolism [39], leading to mitochondrial oxidative phosphorylation and ATP synthesis [25, 40]. Therefore, we postulate that the increased bone formation unique to the SiN material may, in part, be activated by the ammonium released from SiN, supporting the glutamine-driven mitochondrial oxidative phosphorylation system.

Angiogenesis is critical for supplying nutrients and oxygen required for the formation and maturation of bone [41]. Therefore, bone-conductive biomaterials have been designed to induce neovascularization by incorporating VEGF [42, 43] and arginine-glycine-asparate (RGD) adhesion molecule [44] that target vascular endothelial cells [45, 46]. Recent *in vitro* studies have reported that SiN-doped micro-arc oxidation coatings increase human umbilical vein endothelial cell (HUVEC) migration [13] and that silicon oxynitrophosphide (SiOP) coatings increase tube formation and expression of *HIF1a* via HUVEC [47]. The present study analyzed RNA sequence data and revealed upregulated functional gene clusters for vascular system generation in BM-MSC exposed to SiN (**Figure 4**). BM-MSCs are not vascular endothelial cells; however, SiN and its released chemical components may have specific cellular and molecular effects that mimic vascular formation in BM-MSC.

Owing to the cascade of chemical reactions following the release of nitrogen and its subsequent reaction with water molecules, SiN leads to the formation of NO in aqueous solutions [22, 23]. This is known to stimulate angiogenesis [48, 49]. Accordingly, the local environment around the SiN implant may stimulate the endothelial precursor cells to form vascular structures without the use of VEGF and RGD. The present immunohistochemical study identified increased CD31^dim^VEGFR3^+^ vascular-like structures in the bone marrow near the SiN implant (**Figure 8**). CD31 is a vascular endothelial cell marker that is strongly expressed in the type H endothelium of the metaphysis but is less expressed in sinusoidal vessels or the type L endothelium of the diaphysis [50]. VEGFR3 is present in all endothelial cells during development and becomes restricted to the lymphatic endothelium in the adult [51]. However, VEGFR3 is found in the microvasculature of the wound and supports the growth of sprouting new blood vessels from the pre-existing vasculature [52]. Therefore, the CD31^dim^VEGFR3^+^ vascular-like structure was considered a sign of neovascularization induced by the SiN implant.

Osseointegration, measured by the load-resistant push-in test, indicates the strength of bone-implant bonding [21, 29]. SiN and ATi implants were comparable in establishing the BIC (**Figure 7**), whereas the ATi implant exhibited a larger push-in value than the SiN implant (**Figure 4**). This observation may be explained by mechanical interdigitation of the micro-rough surface of the ATi implant, although the interface tissue on the implant surface may be differentially modulated by the micro-rough surface [21] and discrete nanocrystalline surface modification [53]. The present study did not investigate this mechanism further.

Orthopedic and dental implants have been accepted as reliable treatments for long-term survival. However, the clinical success of these elective surgical treatments is influenced by patient selection, with careful exclusion of underlying conditions that might negatively affect outcomes, such as diabetes and severe osteoporosis. Therefore, new and improved materials are needed for clinical applications. The results of the present study suggest that SiN represents a novel implant material that exhibits increased osteoconductive activity through activation of the mitochondrial oxidative phosphorylation system. Furthermore, SiN-associated neovascularization may support the maturation and long-term health of integrated bone tissue.

## 4. Methods

### 4.1. Ethics statement

All animal procedures were approved by the UCLA Animal Research Committee (ARC #1997-136) and followed the PHS Policy for Humane Care and Use of Laboratory Animals. All animals had free access to food and water and were maintained in regular housing with a 12- 12h light/dark cycle.

### 4.2. Experimental materials and surface characterizations

For *in vitro* studies, disk samples (12.7 mm diameter, 1 mm thick) were produced using commercially pure Ti (Grade 2, ASTM F67, Vincent Metals, Minneapolis MN, USA), Zr (TZ-3Y, Tosoh USA, Grove City, OH, USA), and SiN (Si_3_N_4,_ SINTX Technologies, Salt Lake City, UT, USA). Ti disks were produced by parting disks of the desired diameter from a stock rod using a lathe and then machining the surfaces of the disks. Zr and SiN samples were produced by pressing the spray-dried powders into compacts, which were then compacted into disks, firing the disks to remove organic components and densify the ceramic bodies. Zr was sintered at 1400°C for 2 hr in air to greater than 99% theoretical density in accordance with product literature supplied by Tosoh. SiN containing 10% total by mass Al_2_O_3_ and Y_2_O_3_ sintering aids was sintered and hot-isostatically pressed to a theoretical density greater than 99%, as described in a previous study [54].

For *in vivo* studies, miniature implants (0.8 mm diameter, 6 mm long) were produced in commercially pure Ti and SiN (SINTX Technologies, Salt Lake City, UT). The Ti implants were machined directly from a stock wire of the desired diameter. SiN implants were pressed, green- machined, and sintered into blanks as described above for the disk samples. The blanks were then diamond-ground to achieve the desired final geometry.

Ti disks and Ti miniature implants were subjected to the acid-etch protocol using 67% sulfuric acid (Sigma-Aldrich, St. Louis, MO, USA) at 115°C for 75 seconds as previously reported [55]. The untreated Ti material was designated as machined Ti (MTi) and the sulfuric acid-treated Ti material was designated as ATi. All the materials were sterilized under UV light for 30 min on each surface.

The surface morphologies of the materials were examined using scanning electron microscopy (SEM, XL30, Philips, Eindhoven, Netherlands) and laser profile microscopy (VK- 8500, Keyence, Osaka, Japan). The surface topography was evaluated for the average roughness (Sa), peak-to-valley roughness (Sz), and developed interfacial area ratio (Sdr). The wettability of the material surface was determined by measuring the contact angle of 3 µl ddH2O.

### 4.3. BM-MSC

Human BM-MSC (iMSCs: Applied Biological Materials, Richmond, BC, Canada) was expanded in Dulbecco’s modified Eagle’s medium (Life Technologies, Grand Island, NY, USA) with 15% fetal bovine serum (FBS; Life Technologies, Grand Island, NY, USA) and 100 U penicillin/0.1 mg/ml streptomycin (Life Technologies) at 37°C, 5% CO_2_ in a humidified incubator. Human BM-MSC from passages two to eight were used in this study. The cells were seeded on each disk in 24-well culture plates (1 × 10^5^ cells/well) and cultured in osteogenic induction medium consisting of Eagle’s minimum essential medium, alpha modification (MEM α; Life Technologies), supplemented with 15% FBS, 0.1 μM dexamethasone (Sigma-Aldrich, Saint Louis, MO, USA), 10 mM β-glycerophosphate (Sigma-Aldrich), 50 μg/ml ascorbic acid (Sigma-Aldrich), and 250 ng/ml amphotericinB (Thermo Fisher Scientific, Waltham, MA, USA). The culture medium was changed every three days.

After seven days of osteogenic induction, human BM-MSC attached to the disks were transferred to a new 24-well culture plate. The cells were incubated in 10% WST-1 reagent (Thermo Fisher Scientific) in an osteogenic induction medium for 1 h. The supernatant was placed in a 96-well plate and the absorbance was measured at 405 nm using a microplate reader.

After 21 days of osteogenic induction, human BM-MSC were fixed with 10% formalin neutral buffer solution (Thermo Fisher Scientific) and immersed in 40 mM alizarin red S (Sigma-Aldrich) solution (pH 4.1) for 20 min with gentle shaking, followed by washing with distilled water. To quantify the staining results, 10% acetic acid (Sigma-Aldrich) was added to each well and the plate was incubated at room temperature for 30 min with shaking. Loosely attached cells were scraped in acetic acid, transferred to microcentrifuge tubes, and vortexed for 30 s. The samples were heated at 85°C for 10 min and cooled on ice for 5 min. After centrifugation at 20,000 *g* for 15 min, 10% ammonium hydroxide (Thermo Fisher Scientific) was added. Finally, the supernatant was placed in a 96-well plate and the absorbance was measured at 405 nm using a microplate reader.

### 4.4. Osteogenic differentiation RT-PCR

BM-MSC were seeded on SiN, MTi, ATi, and Zr disks in 24-well plates at 40,000 cells/well. After cells became confluent, cells were treated with the osteogenic medium (StemPro® Osteogenesis Differentiation kit (Thermo Fisher Scientific)). The culture medium was replaced twice a week. Total RNA was extracted from cultured BM-MSC on experimental days 7 and 14 using the RNeasy kit (Qiagen, USA) and then analyzed for quality and concentration using a NanoDrop (Thermo Fisher Scientific). cDNA was synthesized using a High-Capacity RNA-to-cDNA kit (Thermo Fisher Scientific), following the manufacturer’s protocol. Quantitative real-time PCR (qPCR) was carried out using TaqMan primer/probe sets (Thermo Fisher Scientific), as per manufacturer instructions, for genes Runx family transcription factor 2 (RUNX2: Hs01047973_m1), alkaline phosphate (ALP: Hs01029142_m1), osteopontin (OPN: Hs00959010_m1), alpha 1 type I collagen (COL1A1: Hs00164004_m1) and osteocalcin (OCN: Hs01587814_g1).

### 4.5. RNA sequencing, bioinformatic analysis and validation RT-PCR

Total RNA samples from BM-MSC cultured on SiN, MTi, and ATi disks for 21 days were extracted using the Quick RNA Miniprep kit (Zymo Research, Inc. Irvine, CA). Total RNA samples from 3 separate wells in each group were combined in equal quantities. RNA-Seq libraries were prepared using a KAPA Stranded mRNA-Seq kit. The workflow consisted of mRNA enrichment and fragmentation, first-strand cDNA synthesis using random priming followed by second-strand synthesis, conversion of the cDNA:RNA hybrid to double-stranded cDNA (dscDNA), and incorporation of dUTP into the second cDNA strand. cDNA was synthesized by end repair to generate blunt ends, A-tailing, adaptor ligation, and PCR amplification. Different adaptors were used to multiplex the samples in one lane. Sequencing was performed in a PE 2x50 run. A data quality check was performed using Illumina SAV. Demultiplexing was performed using Illumina Bcl2fastq v2.19.1.403 software and subjected to RNA sequencing (Illumina NextSeq 500, Illumina, Inc. San Diego, CA, USA). Sequence data were analyzed using the RaNA-seq cloud platform.

Low-quality reads were then removed, data were aligned to the human genome GCHr38, and differential expression was performed using the RaNA-seq cloud platform [56]. The Ensembl transcript release GRCm38.107 GTF were used for the annotation of gene features. For normalization of transcript counts, counts per million (CPM)-normalized counts were generated by adding 1.0E-4. Differential Gene expression was determined via Partek Flow GSA algorithm (Partek® Flow® software, version 7.0 Copyright ©; 2019 Partek Inc., St. Louis, MO, USA). Gene ontology enrichment analysis for differentially expressed genes was performed using ShinyGo 0.77 [57]. The processed data were further analyzed for gene ontogeny of biological pathways, molecular functions, cellular components, and KEGG pathways using gProfiler2 [58].

The expression of GLUL (Hs00365928_g1) and hypoxia-inducible factor 1a (HIF1a: Hs00153153_m1) genes was determined via RT-PCR, as described above.

### 4.6. Mouse femur implant surgery and push-in test

Male 12-week-old C57Bl/6 mice (strain # 000664; RRID: IMSR_JAX:000664; Jackson Laboratory, Bar Harbor, ME) were used. The animals were anesthetized via isoflurane inhalation and the distal femur was exposed. The implant site was prepared using disposable 25-gauge and 23-gauge needles with a rubber stopper placed 6 mm apart and a miniature implant was press-fitted retrogradely, followed by skin closure [29]. Carprofen analgesics (5.0 mg/Kg) (Rimadyl, Zoetis USUS, Parsippany, NJ, USA) was administered subcutaneously at the time of surgery and every 24 h for 2 days after surgery.

After one and three weeks of healing, the mice were euthanized. Implant-containing femurs were harvested and immediately embedded in a test table. The push-in test was performed using a custom-made stainless-steel pushing rod mounted on a 1 kN load cell (Instron, Canton, MA) and the push-in value was determined following a previously described protocol [21].

### 4.7. Micro-CT and bone morphometry

After the push-in test, mouse femurs were recovered and fixed in 10% buffered formalin and applied to micro-CT imaging (SkyScan 1275, Bruker, Billerica, MA, USA) at a source voltage of 60 kV and source current of 166 µA with the scanning resolution of 10 µm per pixel. The ROI for peri-implant bone was defined as 100 µm periphery from the surface along half of the implant (3 mm long) in the bone marrow. Bone morphometric analysis was performed using the CTAn software (Bruker).

### 4.8. Femur bone histology

After micro-CT imaging, the mouse femurs were decalcified with 10% EDTA (Sigma-Aldrich) for 3–4 weeks and longitudinally hemisected along the implant. After the exposed implant was removed, the femur samples were processed for paraffin-embedded sections (4 µm) with hematoxylin and eosin staining.

Photomicrographs were used to measure the BIC using a previously reported method [21]. The thickness of the peri-implant bone was measured using a Java-based image processing program (ImageJ, National Institute of Health, Bethesda, MD, USA).

### 4.9. Immunohistochemistry

The femur sections were subjected to immunohistochemical analyses. The deparaffinized sections were subjected to heat-induced epitope recovery treatment and incubated with primary monoclonal antibodies against CD31 (WM59, Cat# MA1-26196, Thermo Fisher Scientific) and VEGFR3 (AFL4, Cat# 14-5988-82, Thermo Fisher Scientific) at the indicated dilutions, followed by secondary antibody incubation, diaminobenzidine staining, and methylene blue counterstaining. Using VEGFR3 immunohistochemistry photomicrographs, the number of VEGFR3-positive vascular structures was counted.

### 4.10. Statistics

One-way analysis of variance with the post-hoc Tukey-Kramer formula was used to assess differences among multiple experimental groups. Statistical significance was set at p < 0.05.

## Acknowledgements

We thank Dr. Christopher Hamad, Department of Orthopaedic Surgery, UCLA David Geffen School of Medicine, for providing guidance on mouse implant surgery; UCLA Technology Center for Genomics & Bioinformatics for RNA sequencing; and Dr. Yunfeng Li, Translational Pathology Core Laboratory, UCLA David Geffen School of Medicine for immunohistochemistry. This study was supported by a research fund and in-kind materials from SINTX Technologies, Salt Lake City, UT, USA. TK was supported by the JSPS Research Fellowship for Young Scientists (19J117670), SA was supported by the Watanabe Foundation certifying the scholarship Grant Number WS2022-015 and Hiroo and Yukiko NAKATANI Graduate School of Medicine Scholarship Fund 2022. KEK-U was supported by the UCLA QCBio Collaboratory Fellowship awarded by the UCLA Institute for Quantitative and Computational Biosciences (QCB) 2019 – 2024.

## Data Availability

All data generated and analyzed in this study are included in the manuscript. The data will be made available upon request. The RNA sequencing data will be provided by the National Institutes of Health Gene Expression Omnibus at the time of publication.

## Author Contributions

IN: designed the project.

BM, RB: developed the experimental materials.

WG, RK, TO: developed and characterized the experimental materials.

RB, GP: characterized the experimental materials.

WG, RK, IN, AH: performed *in vivo* studies.

TK, SA, TI: performed *in vitro* studies.

IF, AM: isolated human PDL-MSC.

IN, KEK-U: analyzed RNA sequencing data.

IN, AH: supervised the studies.

IN: drafted the initial manuscript.

GP: contributed manuscript and figures.

All authors: contributed to manuscript review

